## Supplementary Information for "Interpretable classification of Alzheimer’s disease pathologies with a convolutional neural network pipeline"

### Supplementary Information for
### Interpretable classification of Alzheimer's disease pathologies with a convolutional neural network pipeline

*Ziqi Tang[1,2], Kangway V. Chuang[1], Charles DeCarli[3], Lee-Way Jin[4], Laurel Beckett[5], Michael J. Keiser[1]*, and Brittany N. Dugger[4]**

[1] Department of Pharmaceutical Chemistry, Department of Bioengineering and Therapeutic Sciences, Institute for Neurodegenerative Diseases, and Bakar Institute for Computational Health Sciences, University of California, San Francisco, San Francisco, CA, USA

[2] School of Pharmaceutical Sciences, Tsinghua University, Beijing, China

[3] Department of Neurology, University of California - Davis School of Medicine, Davis, CA, USA

[4] Department of Pathology and Laboratory Medicine, University of California - Davis School of Medicine, Davis, CA, USA

[5] Department of Public Health Sciences, University of California - Davis, Davis, CA, USA

**Supplementary Table 1** Demographic and neuropathology details of the initial dataset. Abbreviations: AD=Alzheimer's disease; CAA=cerebral amyloid angiopathy; CP=cored plaques; DP=diffuse plaques; hrs=hours; F=Female; M=Male; MTG=middle temporal gyrus; NP=neuritic plaque; PMI=post mortem interval; yrs=years; n/a=not available. For CERAD like scores 0=none, 1=sparse, 2=moderate, 3=frequent.

** As these were archival cases, sections were not available in all regions to determine the exact Thal Amyloid phase (Thal phase).*

| Case # | Group | Age at Death (yrs) | Sex | PMI (hrs) | NIA Reagan | Thal Phase | Braak NFT stage | CERAD NP score | CAA type (1 or 2) | CERAD like Score CP MTG | CERAD like Score DP MTG | CERAD like Score CAA MTG |
|---|---|---|---|---|---|---|---|---|---|---|---|---|
| 1 | train | 81 | M | 7.0 | Not AD | at least Thal 3 | II | 0 | n/a | 1 | 2 | 0 |
| 2 | validation | 74 | M | 19.0 | Low | 3 | II | 0 | n/a | 0 | 1 | 0 |
| 3 | train | 77 | M | 17.0 | Not AD | 1 | II | 1 | n/a | 0 | 1 | 0 |
| 4 | train | 79 | F | 4.6 | Low | 2 | II | 1 | n/a | 0 | 0 | 0 |
| 5 | train | 87 | M | 13.0 | Low | 2 | II | 1 | n/a | 0 | 0 | 0 |
| 6 | train | 77 | F | 2.1 | Low | 3 | III | 1 | 1 | 1 | 2 | 3 |
| 7 | train | 97 | F | 5.1 | Low | 1 | III | 1 | 1 | 1 | 1 | 2 |
| 8 | train | 83 | M | 5.0 | Low | 5 | III | 1 | n/a | 0 | 3 | 0 |
| 9 | train | 80 | F | 3.0 | Intermediate | 4 | IV | 2 | n/a | 1 | 2 | 0 |
| 10 | hold out | 81 | F | 5.0 | High | at least Thal 2 | VI | 2 | n/a | 2 | 2 | 0 |
| 11 | train | 82 | M | 19.0 | High | 4 | VI | 2 | n/a | 2 | 2 | 0 |
| 12 | train | 81 | M | 8.0 | Intermediate | 3 | V | 2 | n/a | 2 | 3 | 0 |
| 13 | train | 83 | M | 9.0 | Intermediate | 5 | IV | 2 | 1 | 0 | 2 | 3 |
| 14 | train | 88 | F | 35.0 | High | 5 | V | 2 | 2 | 2 | 2 | 1 |
| 15 | train | 88 | F | 9.0 | High | 3 | V | 2 | n/a | 0 | 0 | 0 |
| 16 | train | 91 | F | 6.8 | Low | 3 | II | 2 | n/a | 1 | 3 | 0 |
| 17 | train | 89 | M | 8.0 | High | 5 | V | 2 | n/a | 1 | 3 | 0 |
| 18 | validation | 88 | M | 19.0 | High | 4 | VI | 2 | 1 | 2 | 3 | 1 |
| 19 | hold out | 88 | F | 6.2 | Intermediate | 3 | IV | 2 | n/a | 1 | 2 | 0 |
| 20 | train | 62 | M | 7.6 | High | 5 | VI | 2 | n/a | 1 | 3 | 0 |
| 21 | train | 85 | M | 4.5 | High | 5 | VI | 2 | 2 | 1 | 3 | 2 |
| 22 | train | 89 | F | 20.0 | High | 2 | V | 3 | n/a | 0 | 0 | 0 |
| 23 | hold out | 76 | F | 3.3 | High | 4 | V | 3 | n/a | 1 | 2 | 0 |
| 24 | hold out | 78 | F | 7.0 | High | 5 | V | 3 | n/a | 2 | 3 | 0 |
| 25 | hold out | 91 | F | 5.0 | High | 5 | VI | 3 | n/a | 2 | 3 | 0 |
| 26 | train | 85 | F | 4.0 | High | 5 | VI | 3 | n/a | 2 | 2 | 0 |
| 27 | train | 77 | M | 8.0 | High | 5 | V | 3 | n/a | 2 | 3 | 0 |
| 28 | hold out | 81 | M | 4.0 | High | 5 | VI | 3 | n/a | 2 | 3 | 0 |

| | | | | | | | | | | | | |
|---|---|---|---|---|---|---|---|---|---|---|---|---|
| 29 | train | 87 | F | 14.0 | High | 4 | VI | 3 | 1 | 2 | 3 | 1 |
| 30 | train | 62 | F | 4.0 | High | 5 | V | 3 | n/a | 3 | 2 | 0 |
| 31 | train | 83 | M | 2.0 | High | 5 | IV | 3 | 1 | 3 | 2 | 3 |
| 32 | train | 84 | M | 26.0 | High | 5 | V | 3 | 2 | 3 | 2 | 1 |
| 33 | hold out | 83 | F | 4.7 | High | 4 | VI | 3 | 2 | 1 | 3 | 2 |
| 34 | train | 87 | F | 31.0 | High | 5 | VI | 3 | n/a | 2 | 3 | 0 |
| 35 | train | 70 | F | 10.0 | High | 5 | VI | 3 | n/a | 1 | 3 | 0 |
| 36 | validation | 68 | F | n/a | High | 5 | VI | 3 | 1 | 2 | 3 | 1 |
| 37 | validation | 85 | F | 5.0 | Intermediate | 5 | III | 3 | 2 | 2 | 3 | 1 |
| 38 | train | 90 | M | 13.0 | High | 5 | VI | 3 | n/a | 1 | 3 | 0 |
| 39 | hold out | 78 | F | 6.0 | High | 5 | VI | 3 | 2 | 2 | 3 | 1 |
| 40 | hold out | 88 | M | 7.0 | High | 5 | VI | 3 | n/a | 1 | 2 | 0 |
| 41 | hold out | 95 | M | 42.0 | High | 5 | IV | 3 | 2 | 2 | 3 | 1 |
| 42 | train | 92 | F | 4.0 | High | 3 | VI | 3 | 1 | 1 | 2 | 1 |
| 43 | train | n/a | n/a | n/a | n/a | n/a | n/a | n/a | n/a | 2 | 2 | 0 |

**Supplementary Table 2** Demographic and neuropathology details of additional hold out dataset. Abbreviations: AD=Alzheimer's disease; CAA=cerebral amyloid angiopathy; CP=cored plaques; DP=diffuse plaques; hrs=hours; F=Female; M=Male; MTG= middle temporal gyrus; NP=neuritic plaque; PMI=post mortem interval; yrs=years; n/a=not available. For CERAD like scores: 0=none, 1=sparse, 2=moderate, 3=frequent.

| Case # | Group | Age at Death (yrs) | Sex | PMI (hrs) | NIA Reagan | Braak NFT stage | CERAD NP score | CERAD like Score CP MTG | CERAD like Score DP MTG | CERAD like Score CAA MTG |
|---|---|---|---|---|---|---|---|---|---|---|
| 1 | hold out | 83 | M | 26 | Low | II | 0 | 0 | 0 | 0 |
| 2 | hold out | 77 | M | 6.75 | Not AD | I | 0 | 0 | 0 | 0 |
| 3 | hold out | 79 | F | 18 | Low | I | 0 | 1 | 1 | 0 |
| 4 | hold out | 72 | M | 4 | Low | I | 0 | 0 | 1 | 0 |
| 5 | hold out | 93 | F | 5 | Low | II | 1 | 1 | 1 | 0 |
| 6 | hold out | 96 | F | 6.95 | Low | II | 1 | 0 | 3 | 0 |
| 7 | hold out | 78 | F | 9 | Low | III | 1 | 2 | 2 | 3 |
| 8 | hold out | 87 | M | 5 | Intermediate | III | 1 | 0 | 3 | 3 |
| 9 | hold out | 92 | F | 4 | Intermediate | IV | 1 | 2 | 2 | 0 |
| 10 | hold out | 81 | M | 15 | Intermediate | III | 2 | 1 | 1 | 0 |
| 11 | hold out | 81 | M | 7 | High | V | 2 | 1 | 3 | 1 |
| 12 | hold out | 78 | M | 9 | High | V | 3 | 3 | 3 | 0 |
| 13 | hold out | 81 | M | 4 | High | V | 2 | 3 | 3 | 3 |
| 14 | hold out | 81 | F | 20 | Intermediate | IV | 3 | 2 | 2 | 0 |
| 15 | hold out | 90 | F | 76.5 | High | VI | 3 | 0 | 3 | 1 |
| 16 | hold out | 80 | M | 3 | High | VI | 3 | 2 | 3 | 0 |
| 17 | hold out | 89 | M | 7 | High | V | 3 | 2 | 3 | 1 |
| 18 | hold out | 89 | M | n/a | High | VI | 3 | 3 | 3 | 1 |
| 19 | hold out | 69 | M | 4 | High | V | 3 | 2 | 3 | 2 |
| 20 | hold out | 59 | F | 5 | High | VI | 3 | 2 | 3 | 0 |

**Supplementary Table 3** Classification accuracy on validation set and hold-out test set. Using a threshold of 0.91, 0.1 and 0.85 for cored plaque, diffuse plaque, and CAA prediction respectively. Predictions with confidence above the threshold are considered to be positives.

| | Overall | Cored Plaque | Diffuse Plaque | CAA |
|---|---|---|---|---|
| Validation set | 0.970 | 0.978 | 0.938 | 0.993 |
| Hold-out test set | 0.987 | 0.994 | 0.969 | 0.999 |

**Supplementary Table 4** Spearman rank-order correlation coefficient between model-based whole-slide scores and CERAD categories.

| | Cored plaque | Diffuse plaque | CAA |
|---|---|---|---|
| Entire dataset | 0.82 | 0.74 | 0.70 |
| Blinded hold-out set | 0.81 | 0.76 | 0.72 |

### Supplementary Figures

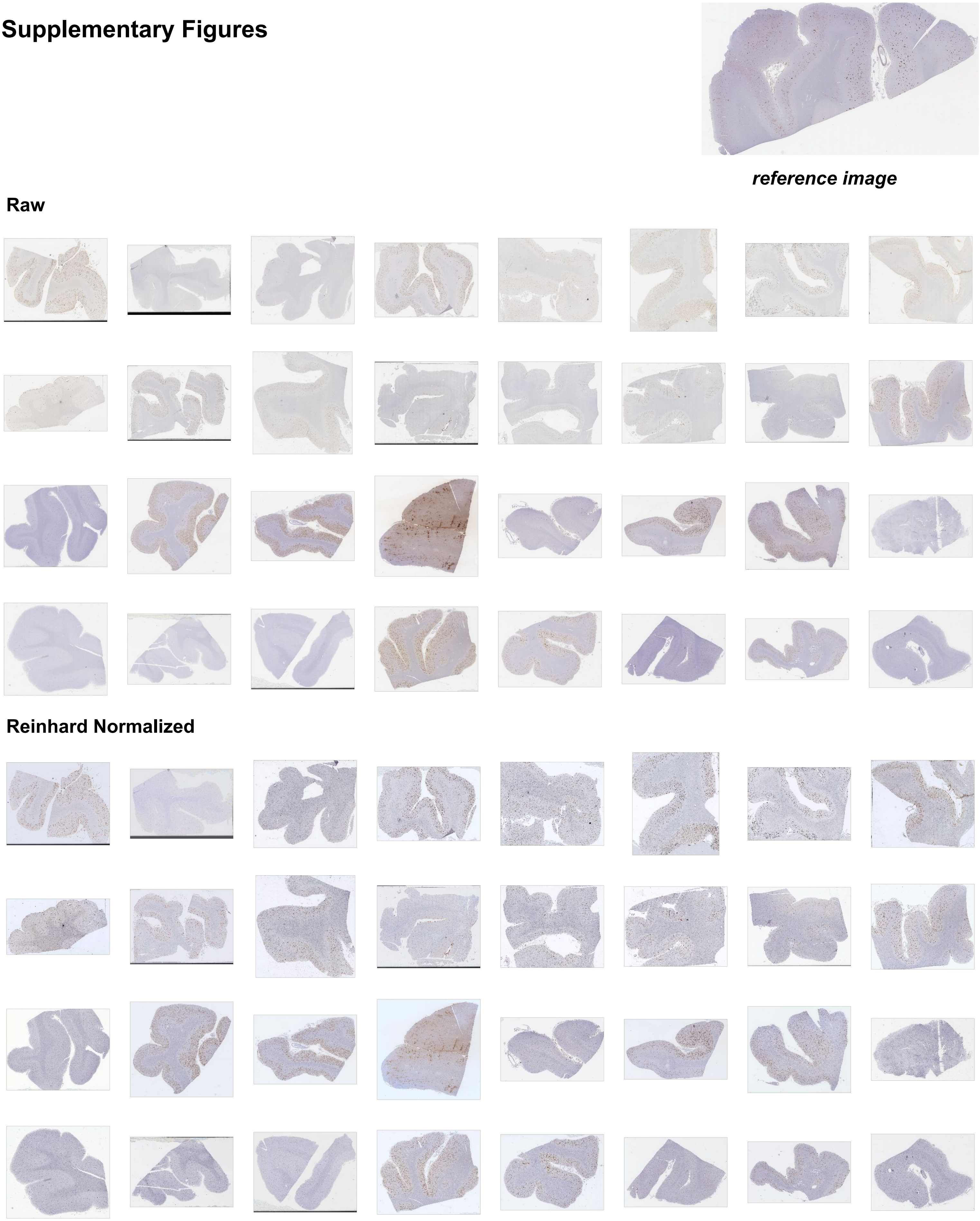


**Supplementary Figure 1** Whole slide image color normalization by the method of Reinhard et. al.

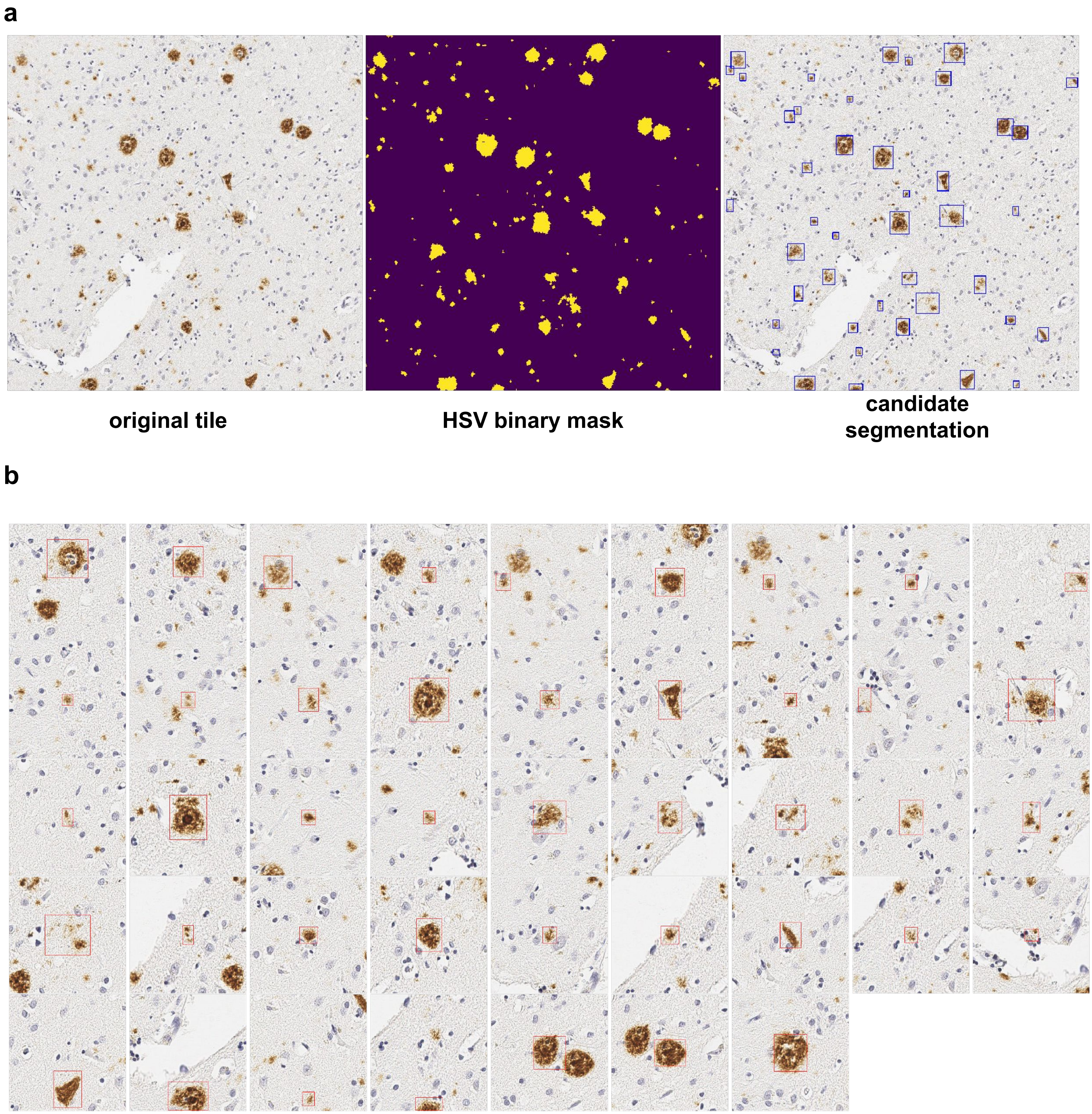


**Supplementary Figure 2** Automatic plaque-candidate segmentation. **a.** A permissive HSV-filter is applied to the original tile and smoothed, providing a binary mask for segmentation. Continuous regions are automatically bounded in minimum boxes and subsequently center cropped. **b.** Resulting center-cropped images with candidate plaques.

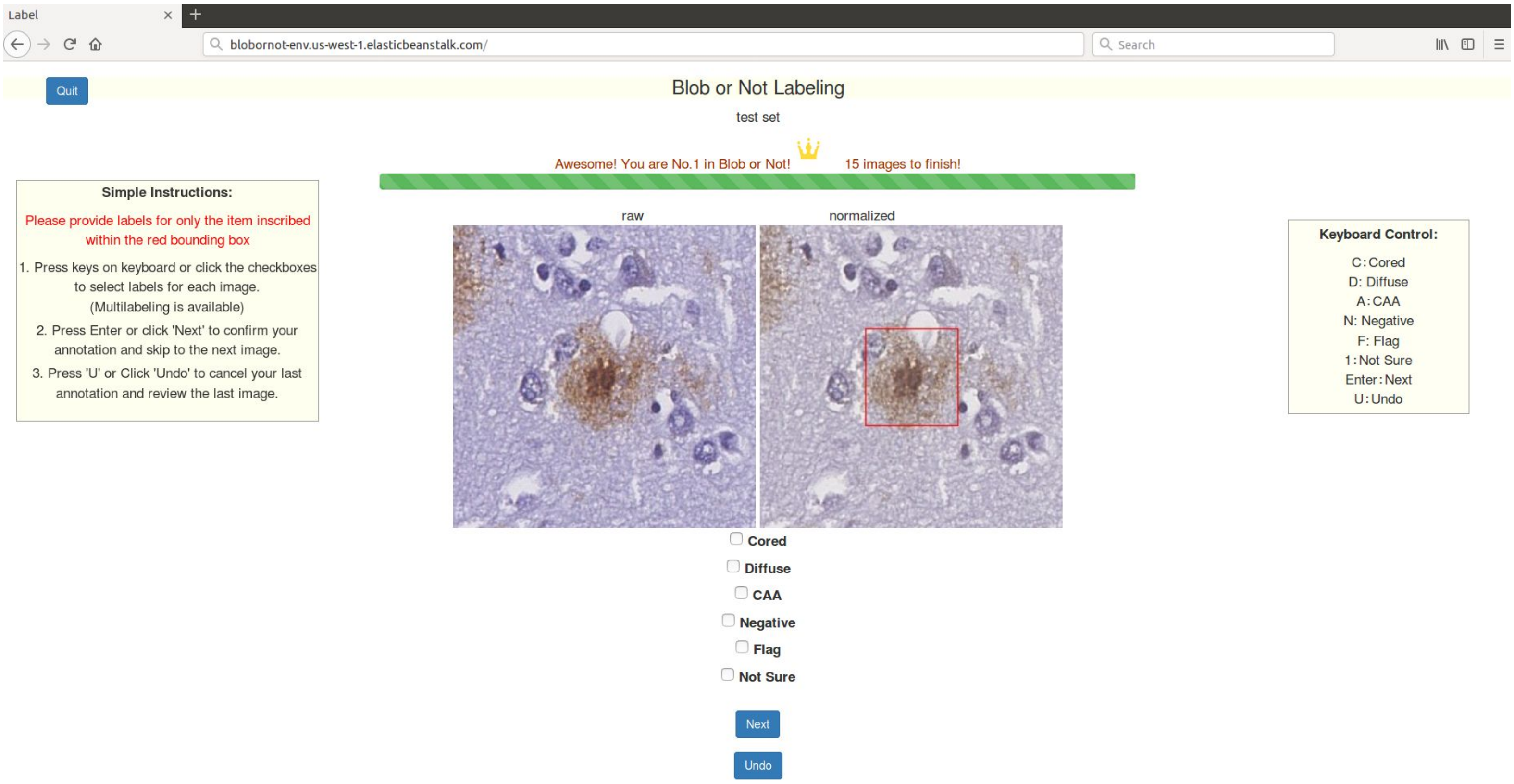


**Supplementary Figure 3** Live screenshot of the custom web platform for rapid image annotation used in this study. Images before and after normalization are shown for comparison. A bounding box in the normalized image specifies which specific blob being labeled. Several elements of gamification, such as leveling, achievement badges (crown icon), and progress bar filling (green bar) are incorporated to reward and motivate annotation task progress. Authorized annotators used a rapid keystroke entry format to label candidate plaques. All labels were stored in a SQL database using Amazon Relational Database Service.

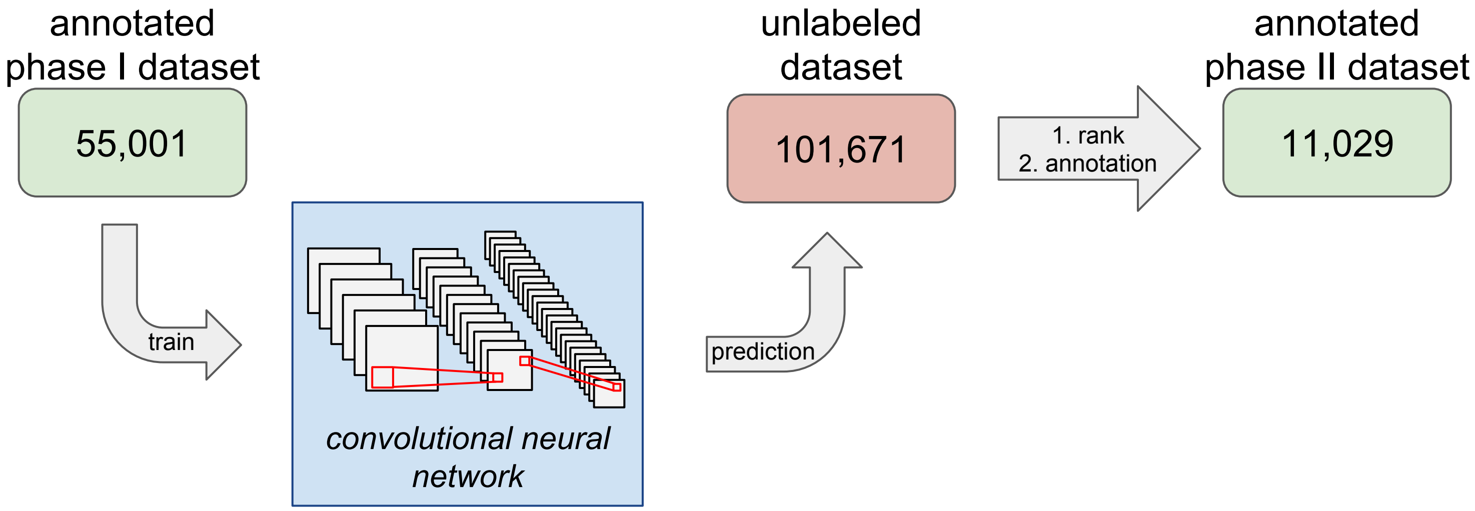


**Supplementary Figure 4** Annotation phase II for minor classes enrichment. An intermediate CNN model for plaques classification was trained on the phase I dataset containing 55,001 annotated images. The trained model was applied to another dataset with 101,671 unlabeled images. These unlabeled images were ranked by the prediction confidence for cored plaques or CAAs. 11,029 images with high prediction confidence for cored plaques or CAAs were then labeled by the neuropathologist as annotation phase II.

**a**

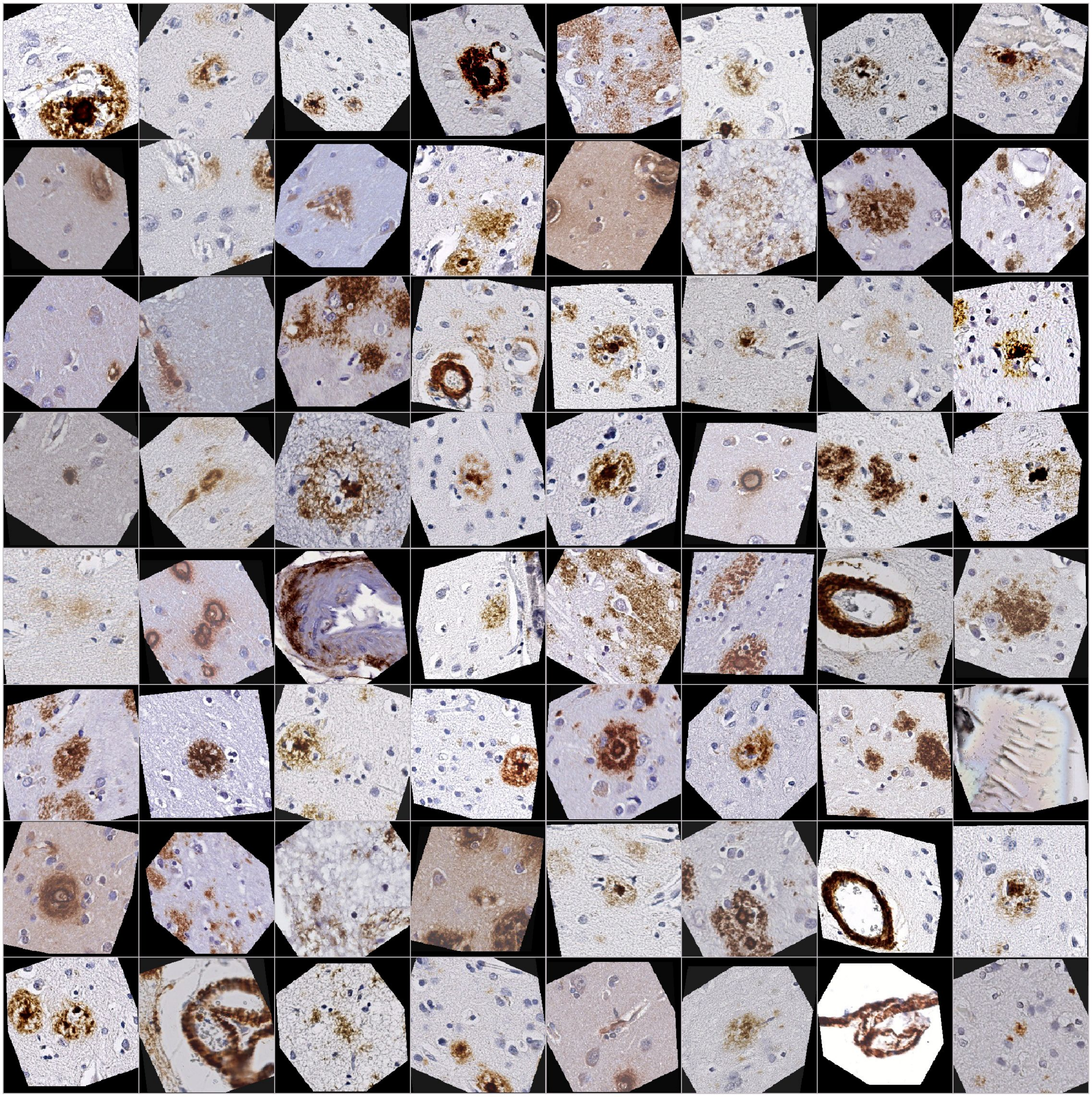

**b**

| Dataset | Cored | Diffuse | CAA | Negative | Total |
|---|---|---|---|---|---|
| **Raw** | 2,141 (3.49 %) | 48,123 (78.41%) | 2,227 (3.63%) | 3462 (5.64%) | 61,370 |
| **Upsampling** | 49,411 (31.83 %) | 67,008 (43.16 %) | 49,170 (31.67 %) | 3462 (2.23%) | 155,239 |

**Supplementary Figure 5** CNN training was performed with real time augmentation and minority class upsampling. **a.** Training examples shown with data-augmentation, including random flips, rotations, zooms, and color jitter. **b.** Training was evaluated with and without minority-class upsampling.

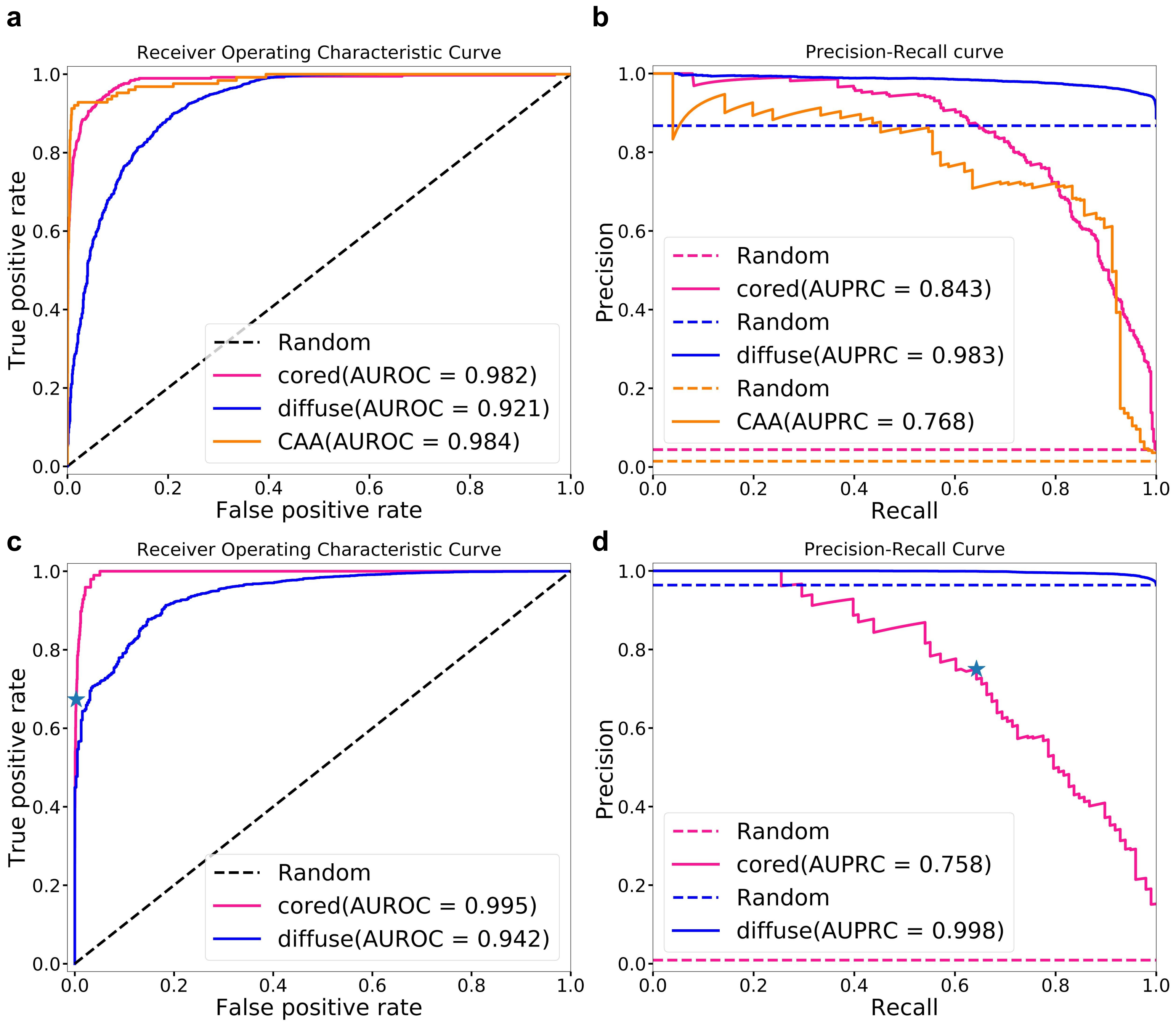


**Supplementary Figure 6** Prediction performance without color normalization. **a**. Receiver operator characteristic curve (on validation set). **b.** Precision-recall curve (on validation set). **c**. Receiver operator characteristic curve (on hold-out set). **d.** Precision-recall curve (on hold-out set). Source data are provided as a Source Date file.

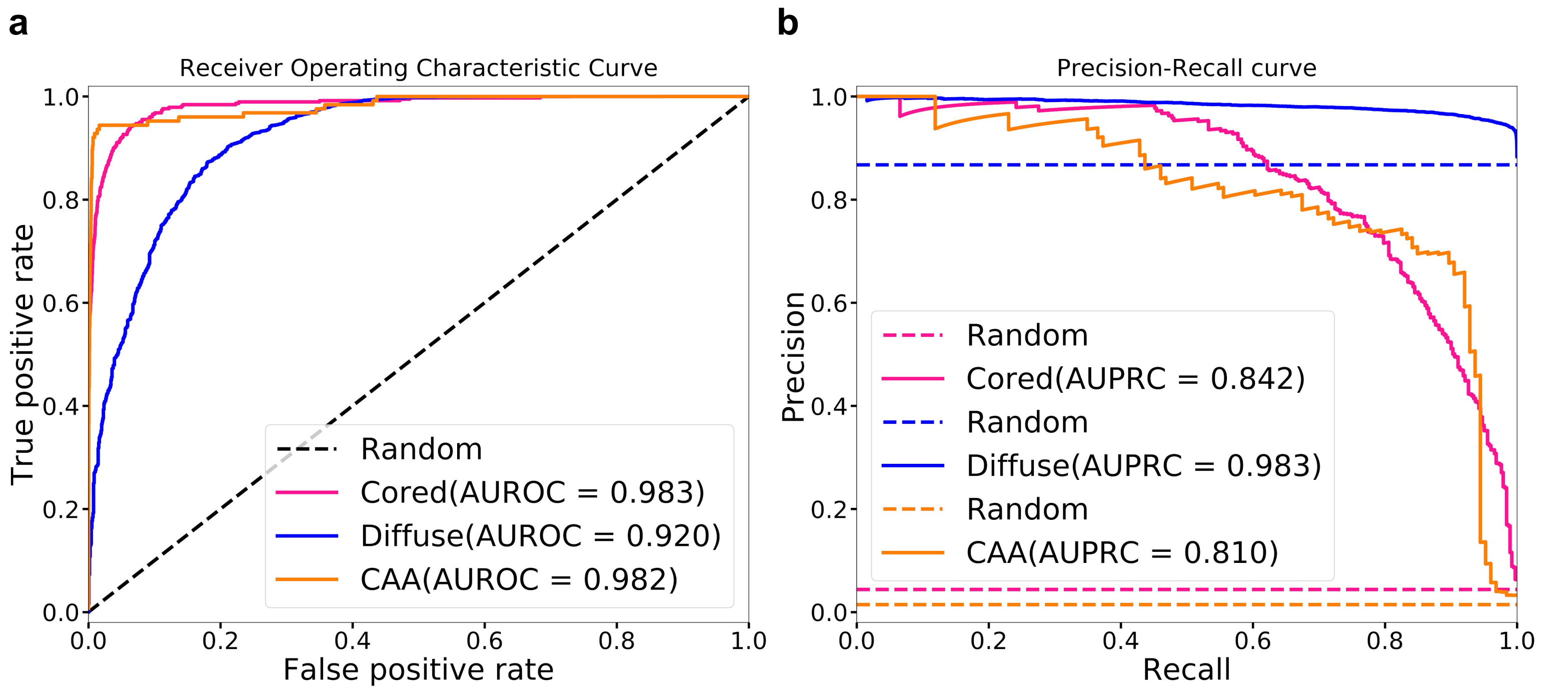


**Supplementary Figure 7** Prediction performance on validation set. **a**. Receiver operator characteristic curve. **b.** Precision-recall curve. Source data are provided as a Source Date file.

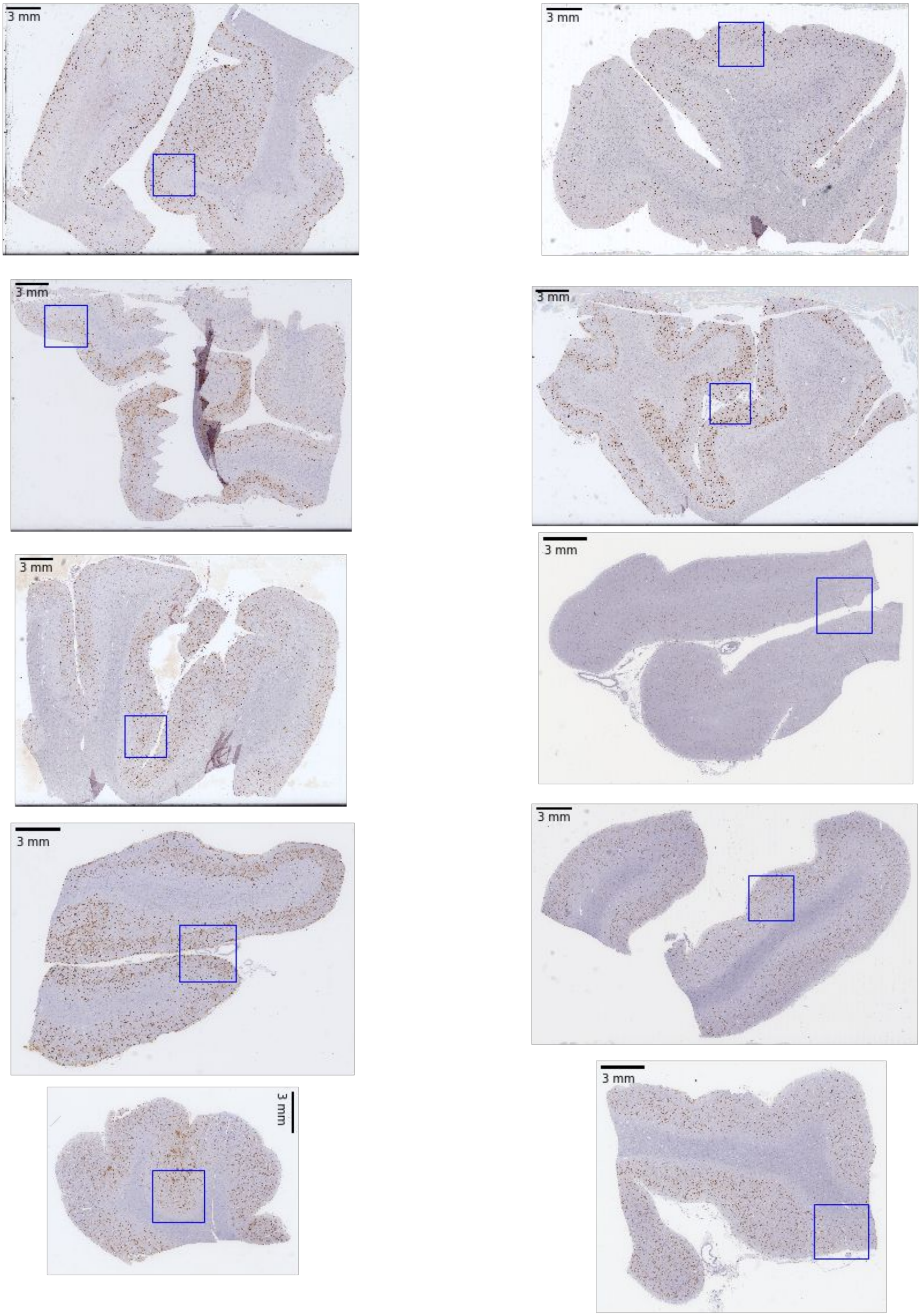


**Supplementary Figure 8** Hold-out test set WSIs. Continuous regions of interest (corresponding to 3.8 x 3.8 mm tissue regions) are outlined in blue. Candidate plaques were extracted using the segmentation script for local analysis.

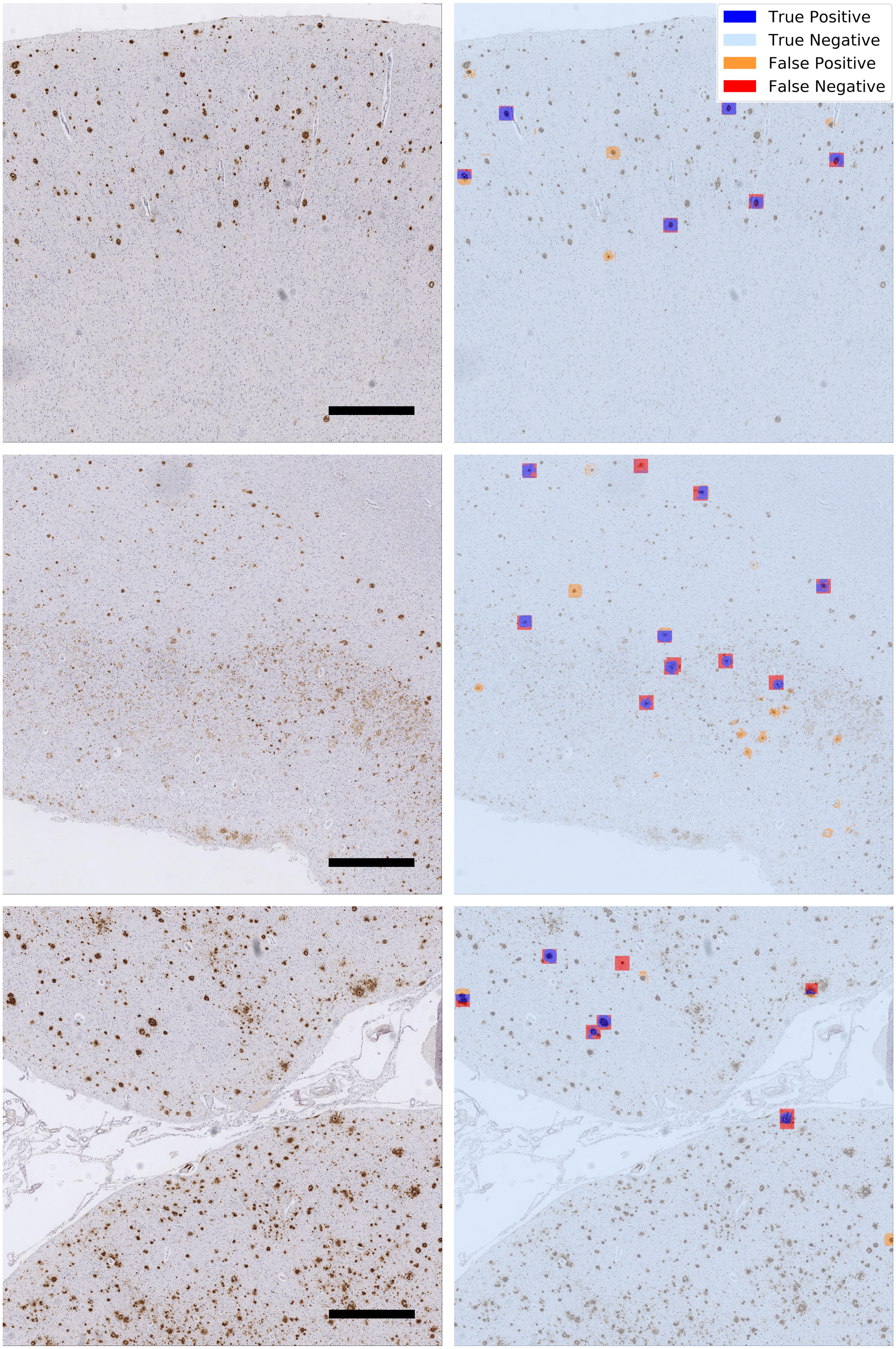


**Supplementary Figure 9** Additional examples of visual representation of classification performance on held-out set against ground truth plaque labels. Overlaying prediction confidence maps with expert labeled cored plaque bounding-boxes provides a color-coded method for visualizing network performance. The resulting plot (right column) shows the assessment of agreement between CNN predictions and expert labels, in terms of true positives (blue), false positives (orange), true negatives (cyan), and false negatives (red). Scale bar=750 um.

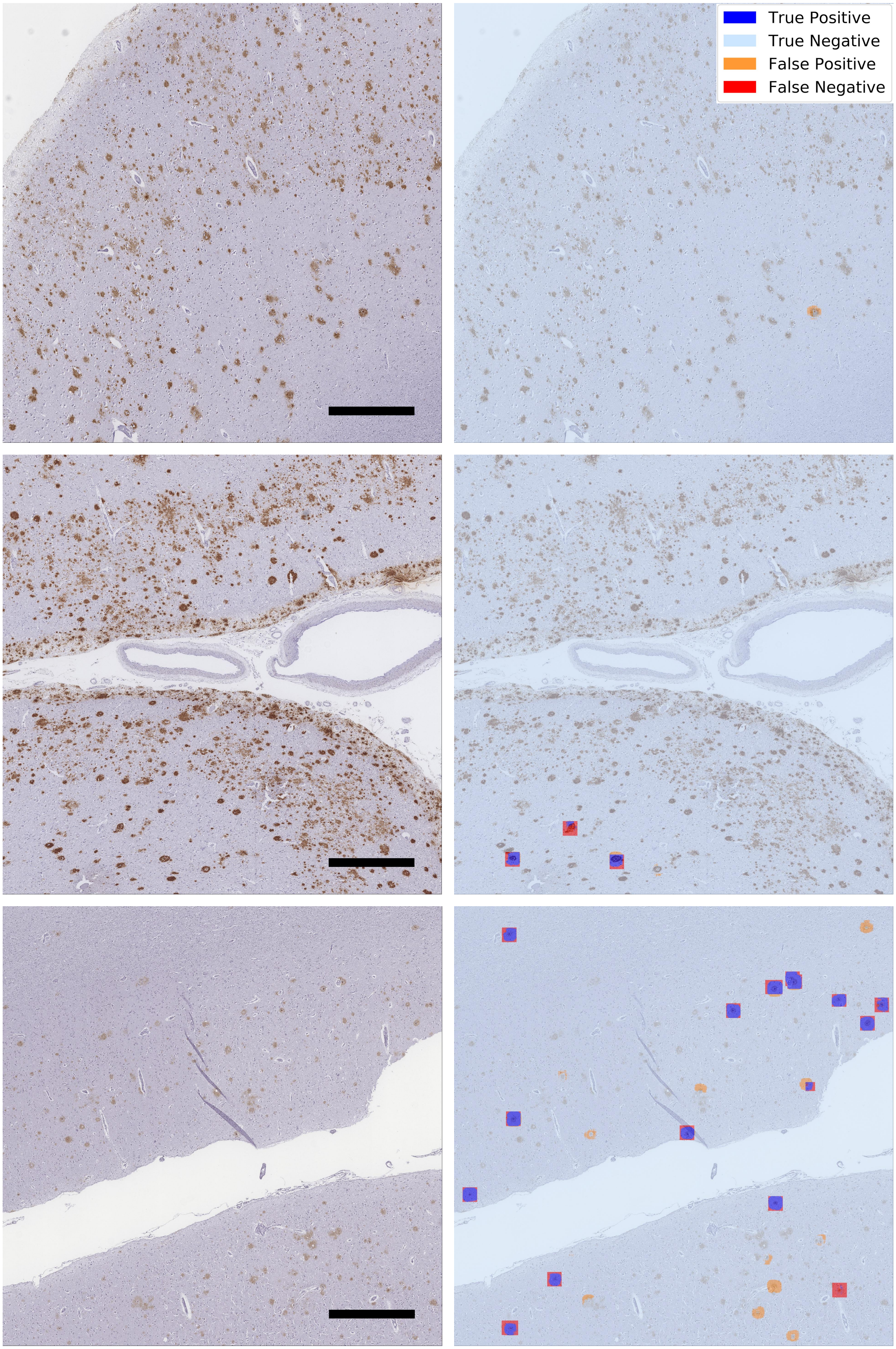


**Supplementary Figure 9 continued**

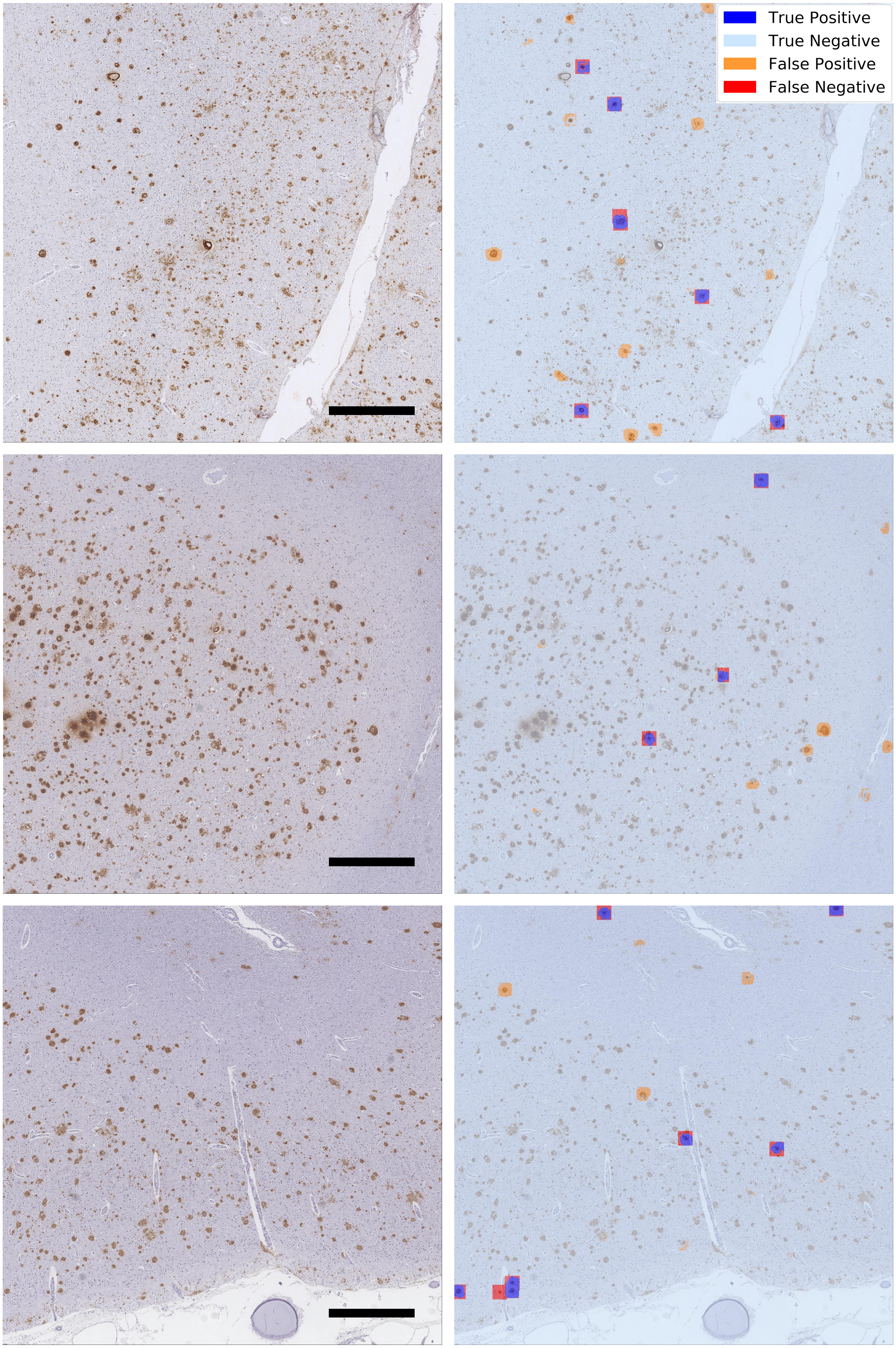


**Supplementary Figure 9 continued**

a

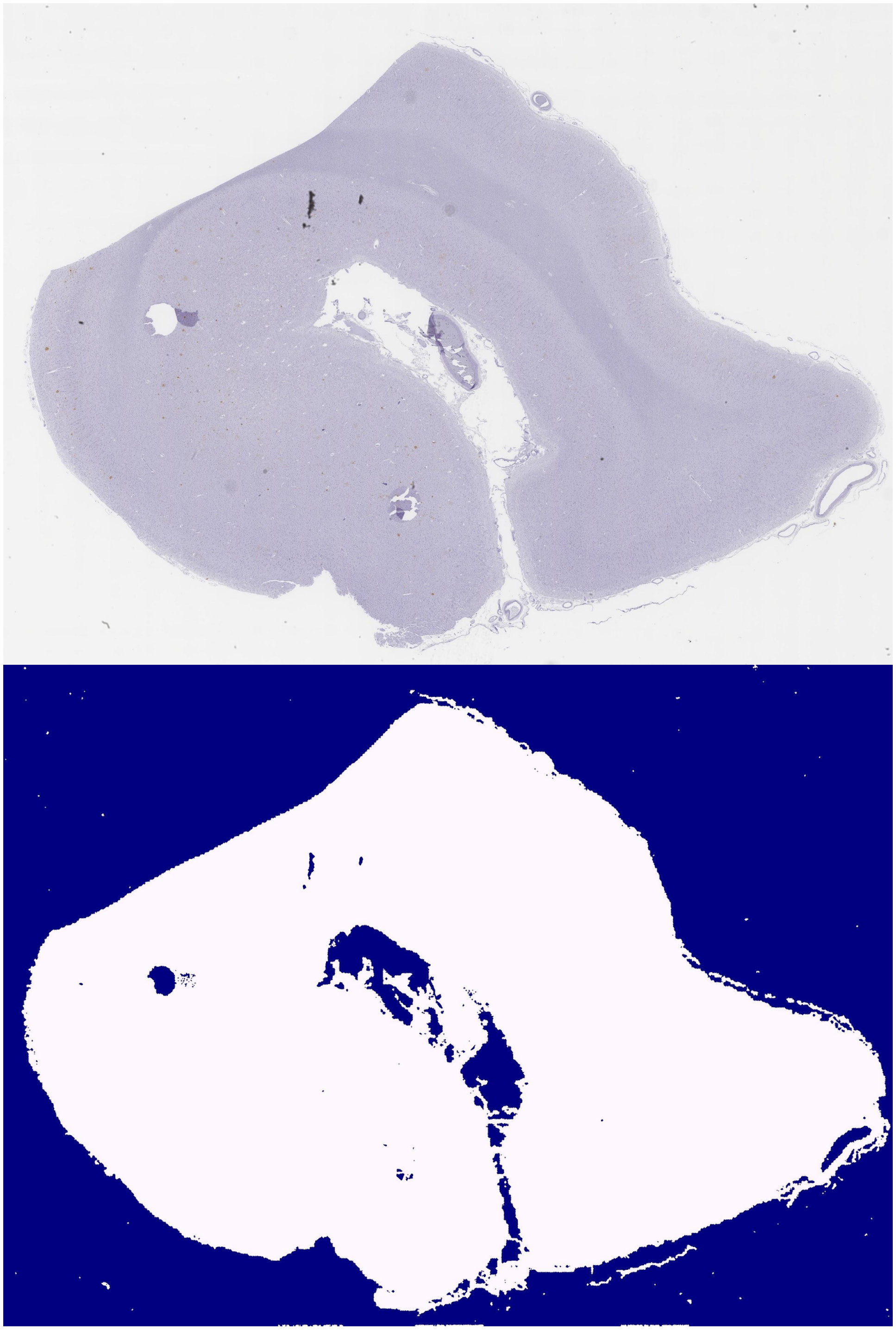

b

**Supplementary Figure 10** Tissue segmentation example. Tissue segmentation was performed in the lightness-chroma-hue (LCH) colorspace utilizing a specific colormask for each WSI. Morphological opening and closing operations were performed to smooth the colormasks, and the tissue areas were calculated as the pixel sum of the refined mask. **a.** IHC stained whole slide image. **b.** Segmented mask.

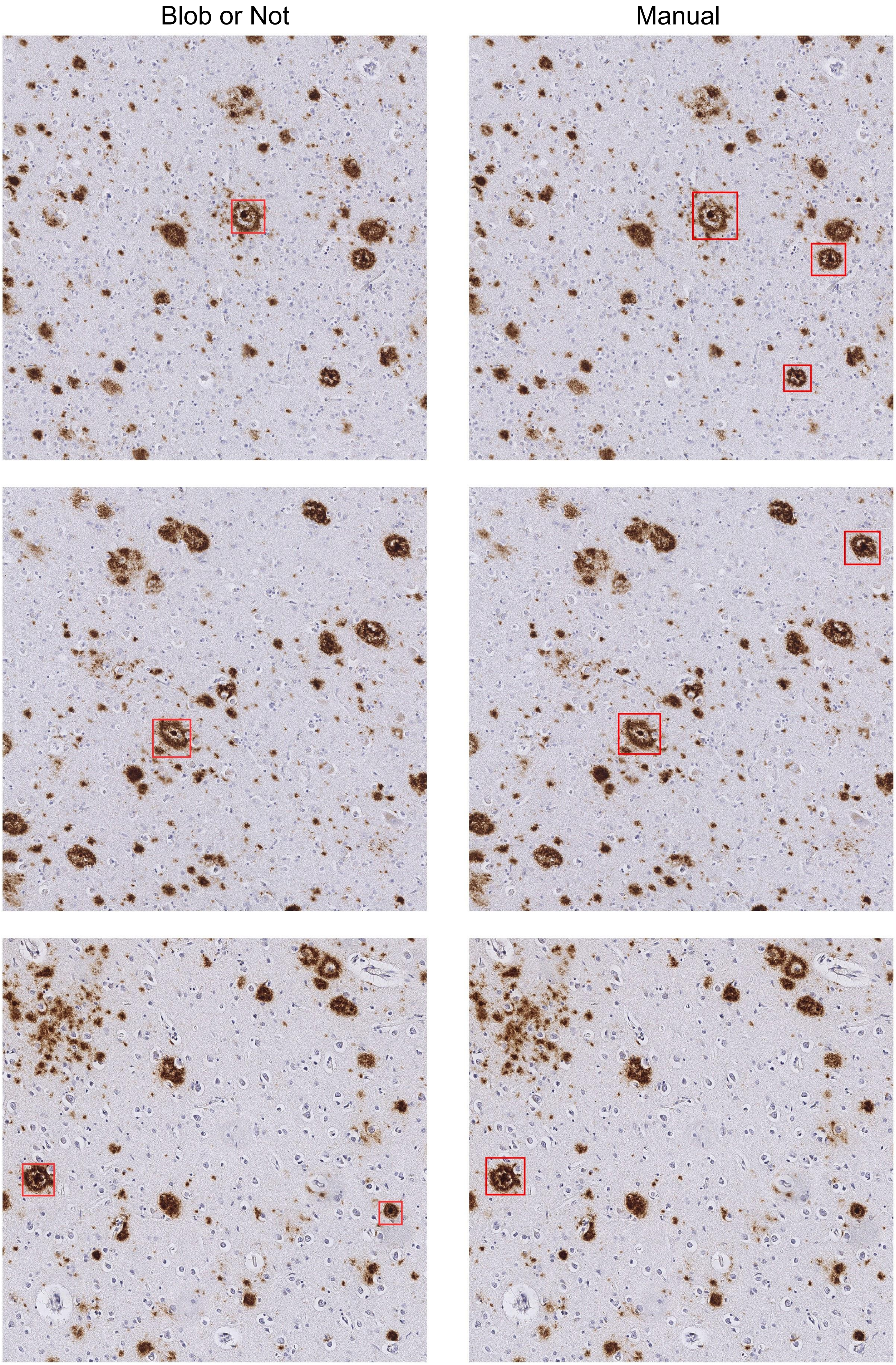


**Supplementary Figure 11** Label comparison. Bounding boxes mark cored-plaque expert annotations

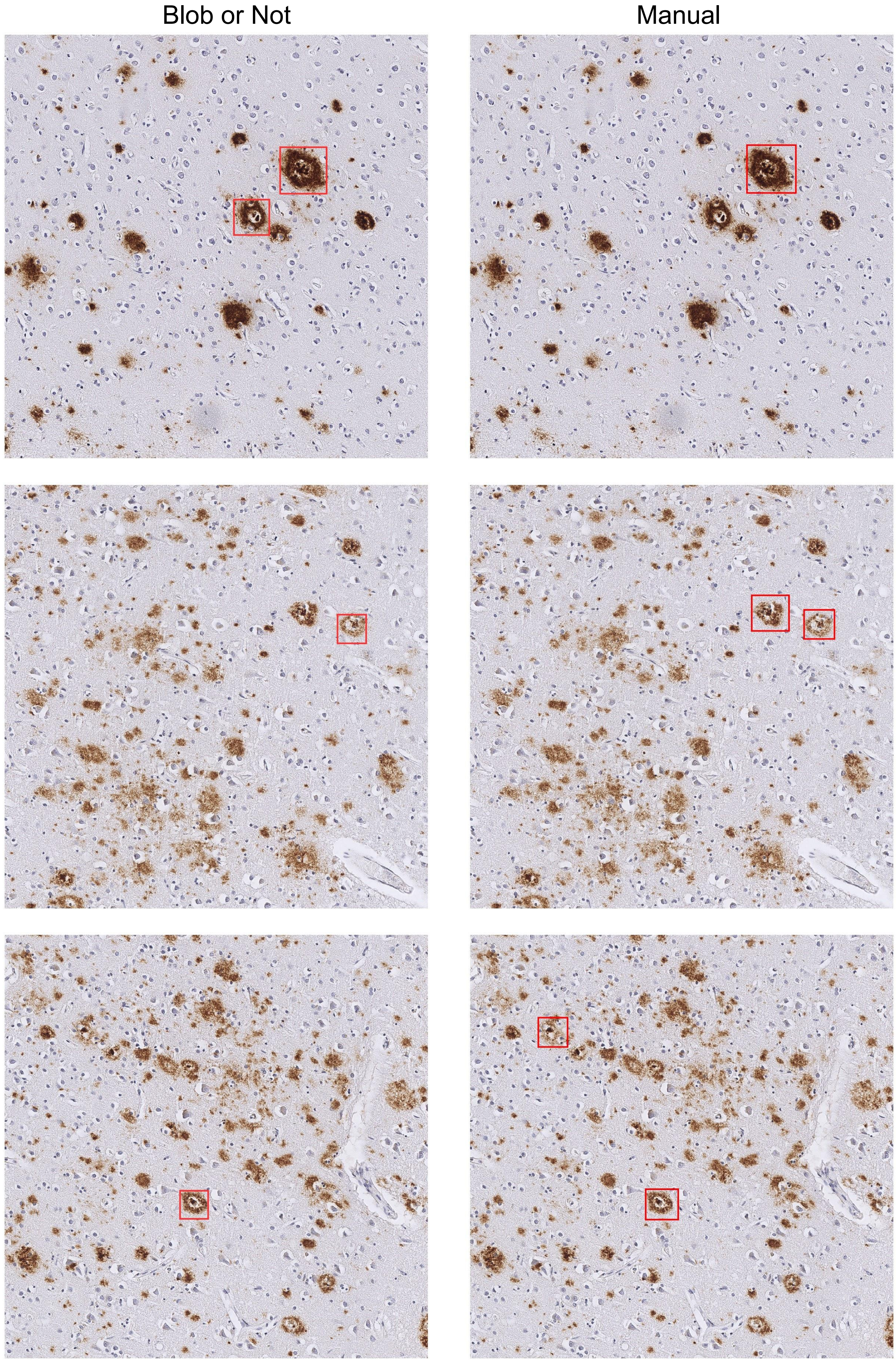


**Supplementary Figure 11 continued**

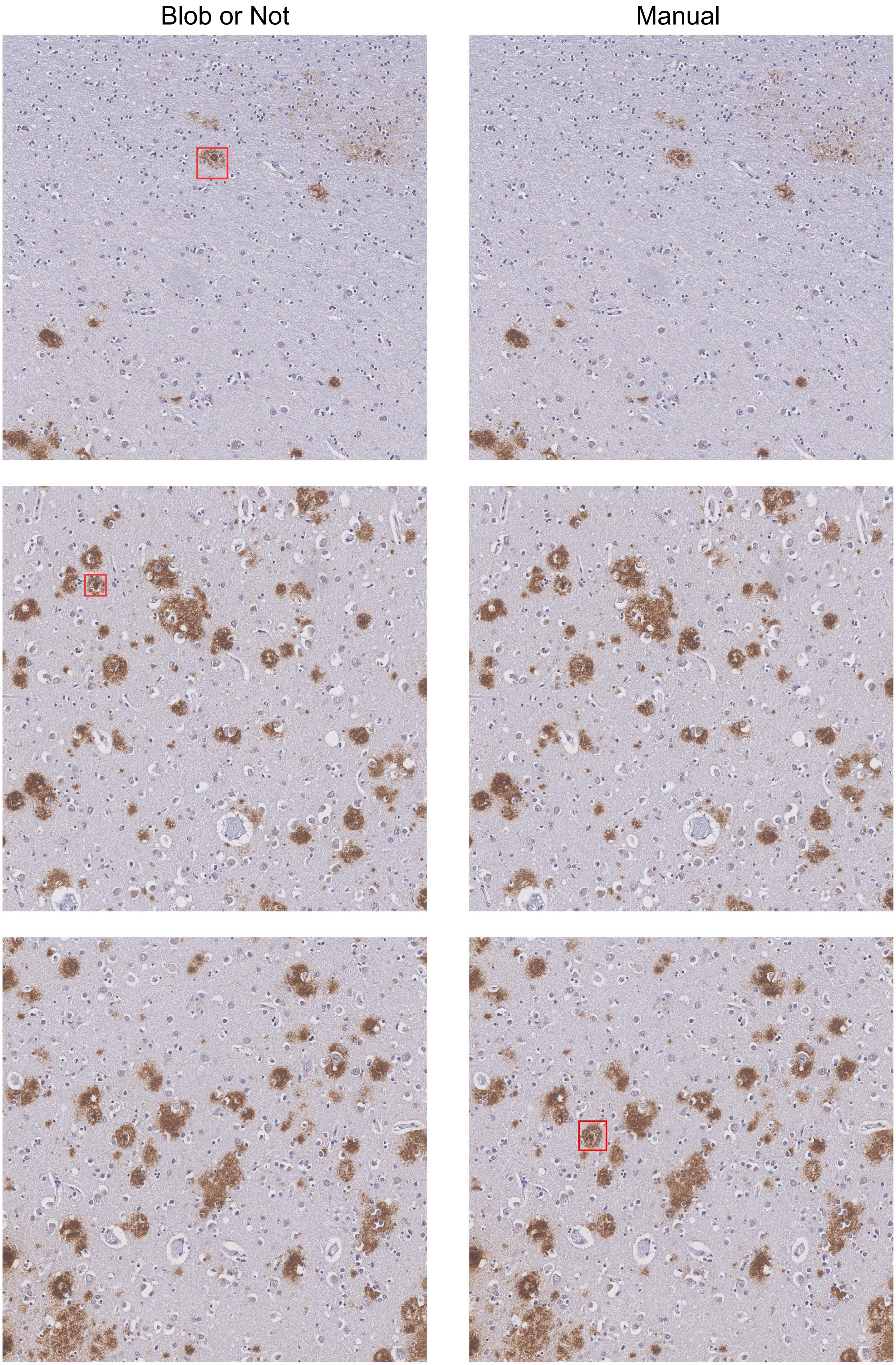


**Supplementary Figure 11 continued**
